# CO_2_ supply modulates lipid remodelling, photosynthetic and respiratory activities in *Chlorella* species

**DOI:** 10.1101/2021.02.18.431842

**Authors:** Michela Cecchin, Matteo Paloschi, Giovanni Busnardo, Stefano Cazzaniga, Stephan Cuine, Yonghua Li-Beisson, Lutz Wobbe, Matteo Ballottari

## Abstract

Microalgae represent potential solutions to reduce the atmospheric CO_2_ level through photosynthesis. To boost CO_2_ fixation by microalgae it is essential to understand physiologic and metabolic responses at the base of CO_2_ assimilation and carbon flow. In this work two *Trebouxiophyceae* species, *Chlorella sorokiniana* and *Chlorella vulgaris*, were investigated for their metabolic responses to high and low CO_2_ (air level) availability. High CO_2_ availability resulted in an increase in biomass accumulation in both species but with a different chloroplast and mitochondrial responses. In *C. sorokiniana* we observed increased polar lipids and protein amount and a balanced NADPH redox state and a similar total respiration in the two conditions analysed. In contrast, in *C. vulgaris* high CO_2_ level caused an increase in TAG accumulation and a higher NADPH consumption suggesting a CO_2_ dependent increase of reducing power consumption in the chloroplast, which in turn influences the redox state of the mitochondria by lowering total dark respiration. Several rearrangements of the photosynthetic machinery were observed in both species, which differ from those described for the model organism *Chlamydomonas reinhardtii*. In the case of *C. reinhardtii*, adaptation of the photosynthetic apparatus to different CO_2_ availability relies on the translational repressor NAB1. NAB1 homologous protein could be identified only in *C. vulgaris* but lacked the regulation mechanisms previously described in *C. reinhardtii*. These findings highlight that the acclimation strategies to cope with a fluctuating inorganic carbon supply are diverse among green microalgae and point to new biotechnological strategies to boost CO_2_ fixation.

**One sentence summary:** High/low CO_2_ availability induces cell responses as lipids remodelling, adaptations of the photosynthetic apparatus and modulation of mitochondrial respiration not conserved among green algae

## Introduction

Microalgae emit half of the oxygen available in the atmosphere and contribute to half of the total organic carbon produced worldwide (Li-Beisson et al., 2019; Salomé and Merchant, 2019). Thanks to the photosynthetic process algae convert light energy into chemical energy to fix CO_2_ in organic compounds. Carbon dioxide is one of the main greenhouse gasses responsible for global warming. CO_2_ level at the Earth’s surface atmosphere is constantly increasing reaching 407.4 ± 0.1 ppm for 2018, an increase of 2.4 ± 0.1 ppm from 2017 (Dlugokencky, 2019). There is an urgent need for an efficient way to reduce the global carbon footprint, which is fundamental to reduce the effects of human activity in the worldwide poise.

Microalgae are emerging as a possible solution due to their ability to grow at high levels of CO_2_ and to produce biomass that can be exploited for several applications: as food or feed supplement, biofuels or to produce high value products. Moreover, these photosynthetic organisms do not require arable land, have a fast growth rate and waste products as well as wastewater-derived effluent can be used as fertilizers for their cultivation (Lum et al., 2013).

Light is harvested in the microalgal chloroplast by pigment binding protein complexes called Photosystem I (PSI) and II (PSII). These complexes are composed of a core complex, where photochemical reactions occur, and an external antenna system which increases light harvesting efficiency and where several photoprotective reactions occur (Gao et al., 2018; Pan et al., 2019). In oxygenic photosynthetic organisms, as eukaryotic microalgae, PSI and PSII work in series to strip electrons from water and transfer them to NADP^+^ producing NADPH. During this linear electron transport protons are pumped from stroma to the lumen generating an electrochemical gradient used by ATPase to synthetize ATP. ATP and NADPH are then used by the Calvin Benson cycle to fix CO_2_ into sugars. In parallel, another electron transport chain takes place in mitochondria, consuming oxygen and NADH and releasing NAD^+^ and ATP. A constant balance between chloroplast and mitochondrial activity is fundamental for cell survival and for adaptation to fluctuating environmental conditions.

It is important to point out that CO_2_ diffusion in the water environments, where microalgae live, is strongly reduced compared to CO_2_ diffusion in air. CO_2_ -limitation is known to reduce the consumption of ATP and NADPH by the Calvin Benson cycle leading to an over-reduced photosynthetic electron transport chain, which could potentially lead to oxidative stress (Wang et al., 2015). For this reason several microalgae species evolved an efficient system to enrich the CO_2_ level inside the cell, i.e. Carbon Concentrating Mechanism (CCM), a complex mechanism by which inorganic carbon is actively transported close to the enzyme responsible for its fixation, i.e. the RUBISCO enzyme (Wang et al., 2015). The CCM mechanism is induced by low CO_2_ concentrations (air level or lower) (Wang et al., 2015). CO_2_ availability plays thus a critical role in modulating photosynthetic efficiency and biomass accumulation in microalgal cultures. For example, in the model green alga *Chlamydomonas reinhardtii*, CO_2_ has been reported to act as a molecular switch inducing a complex network of cell adaptation mechanisms including a translational up-regulation in the formation of PSII antenna complexes (Mussgnug et al., 2005; Wobbe et al., 2009; Berger et al., 2014; Berger et al., 2016; Blifernez-Klassen et al., 2021). In conditions of low CO_2_ availability, accumulation of the cytosolic RNA-binding protein NAB1 is triggered by the transcription factor LCRF (Low Carbon dioxide Response Factor) (Blifernez-Klassen et al., 2021). NAB1 then represses the translation of transcripts encoding light-harvesting antenna proteins (Mussgnug et al., 2005; Berger et al., 2014). The translation repressor activity of NAB1 is controlled by two independent mechanisms related to the methylation of Arg90 and Arg92 residues (Blifernez et al., 2011) and to the redox state of Cys181 and Cys226 residues (Wobbe et al., 2009). NAB1 is highly active in the methylated state, while reduced Cys181 and reduced Cys226 are required for NAB1 RNA-binding activity. Nitrosylation of Cys181 and Cys226 has also been reported to inhibit the RNA binding activity of NAB1 (Berger et al., 2016). The truncation of the PSII antenna reduces the excitation pressure on the photosynthetic apparatus as a response to diminished CO_2_ availability (Berger et al., 2014). Accumulation of NAB1 and its post-translational regulation have been demonstrated to be finely tuned as an acclimation mechanism to different environmental conditions, including varying CO_2_ availability (Wobbe et al., 2009; Berger et al., 2014; Berger et al., 2016).

Among microalgae species discovered, *Trebouxiophyceae* represent an evolutionary defined class of green algae (*Chlorophyta*) comprising the green freshwater algae of the *Chlorella* genus, one of the first microalgae to be cultured on a large scale due to their easy cultivation and high resistance to stresses (Yang et al. 2016; Borowitzka et al. 2018). Species belonging to the *Trebouxiophyceae* class are evolutionarily separated from the model species of green alga, *C. reinhardtii*, belonging to the *Chlorophyceae* class. *Chlorella* species are interesting for industrial cultivation, being reported to rapidly accumulate biomass containing high lipid, protein, carotenoid and vitamin amounts (Li et al., 2013; Lum et al., 2013; Treves et al., 2013; Sarayloo et al., 2017; Camacho et al., 2019; Cecchin et al., 2019). However, the lack of genetic resources and the low efficiency of transformation methods has limited the development of genetic engineering in these species (Cecchin et al., 2019; Lin et al., 2019).

In this work two *Chlorella* species, *Chlorella sorokiniana* and *Chlorella vulgaris*, were investigated for their physiologic and metabolic responses to a fluctuating CO_2_ availability. Our results highlighted contrasting metabolic responses to CO_2_ availability among green algae.

## RESULTS

### Species of the genus *Chlorella* adjust their carbon flow differentially in response to high carbon dioxide availability

*C. sorokiniana* and *C. vulgaris* cells were grown in 80 ml batch airlift photobioreactors bubbled with air (CO_2_ concentration of □0.04%, defined as AIR condition throughout the manuscript) or air enriched with 3% of CO_2_ (defined as CO_2_ condition throughout the manuscript). *Chlorella* strains were cultivated at 300 μmol photons m^-2^ s^-1^ until the saturation phase was reached. Growth kinetics were followed by measuring the optical density (OD) at 720 nm and fitted with sigmoidal function as showed in Figure 1. In both species the increased CO_2_ concentration induced a faster growth rate, as highlighted by the first derivative of the growth curves (Figure 1B-1E). Total biomass production was increased in CO_2_ compared to AIR condition by 253% ± 52% in *C. sorokiniana* and 269% ± 13% in *C. vulgaris* (Figure 1C-1F). Moreover, ∼4-fold increase in maximal daily productivities was observed for both species in CO_2_ condition. These data confirmed that in the cultivation conditions applied carbon fixation processes were stimulated in *Chlorella* species by high CO_2_ suggesting a possible carbon-limitation in AIR condition.

Biomass composition at the end of the growth curves was evaluated, and this revealed a significant increase in total lipids in both species, while different effects were observed on protein and starch content per dry weight (Figure 2). Indeed, CO_2_ condition triggered an accumulation of proteins in *C. sorokiniana* with the amount of starch not being altered. In *C. vulgaris*, protein amounts remained stable, but a decrease in starch content was observed. This suggests a different behaviour of the two species in macromolecule accumulation upon increased carbon availability underlying potential differences in metabolic rearrangement.

**Figure 1.**
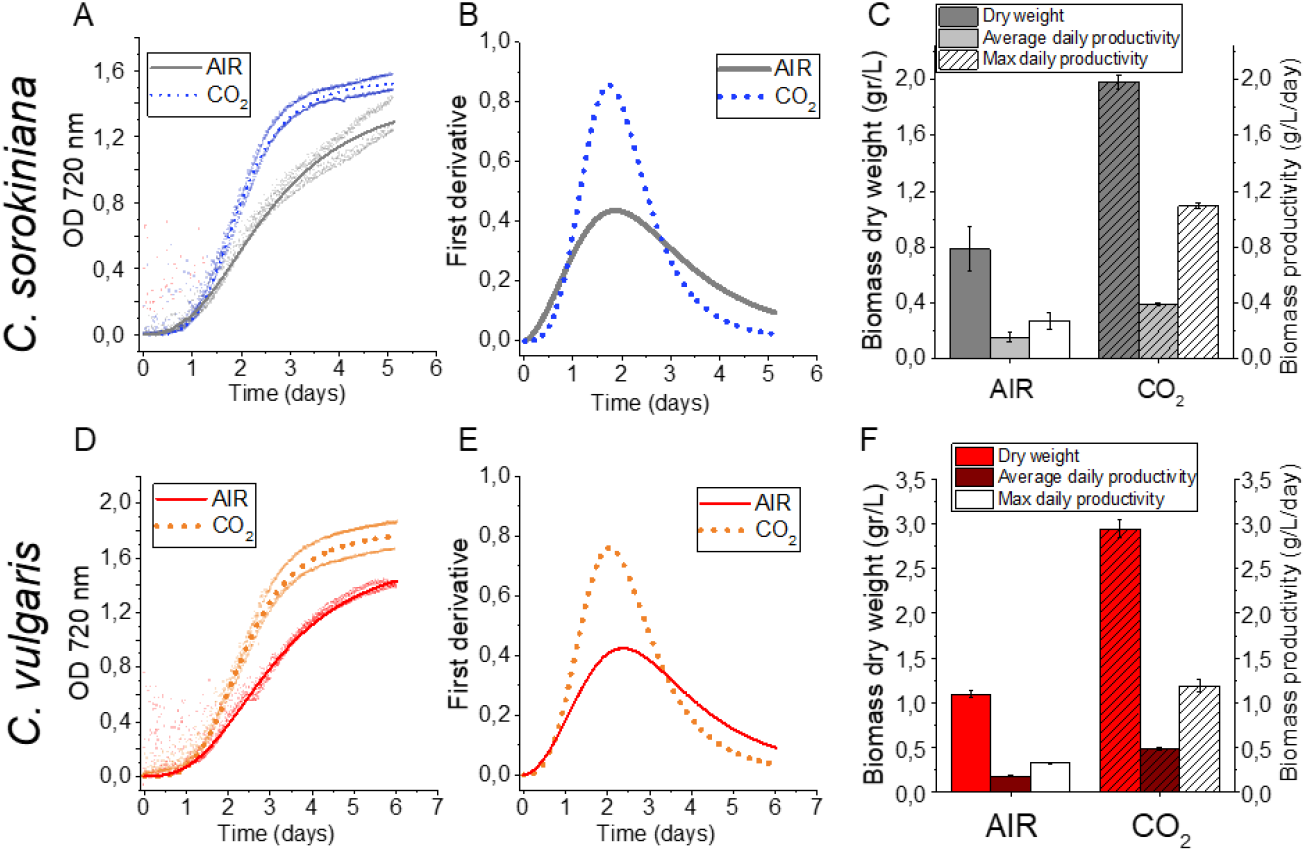
Growth curve and biomass productivity in AIR *vs*. CO_2_. Growth curve and biomass productivity are reported for *C. sorokiniana* in panel A-C and for *C. vulgaris* in panel D-F in AIR condition (□0.04% CO_2_) compared to CO_2_ condition (3% CO_2_). (A, D): growth curve obtained measuring OD at 720nm fitted with sigmoidal function. (B, E): first derivate of growth curves reported in panels A and D. (C, F) Dry weight (g/L), average and maximum daily productivity (g L^-1^ day^-1^) obtained harvesting the biomass at the end of the growth curve. Data are means of 4 replicates and error bars represent standard deviation.

**Figure 2.**
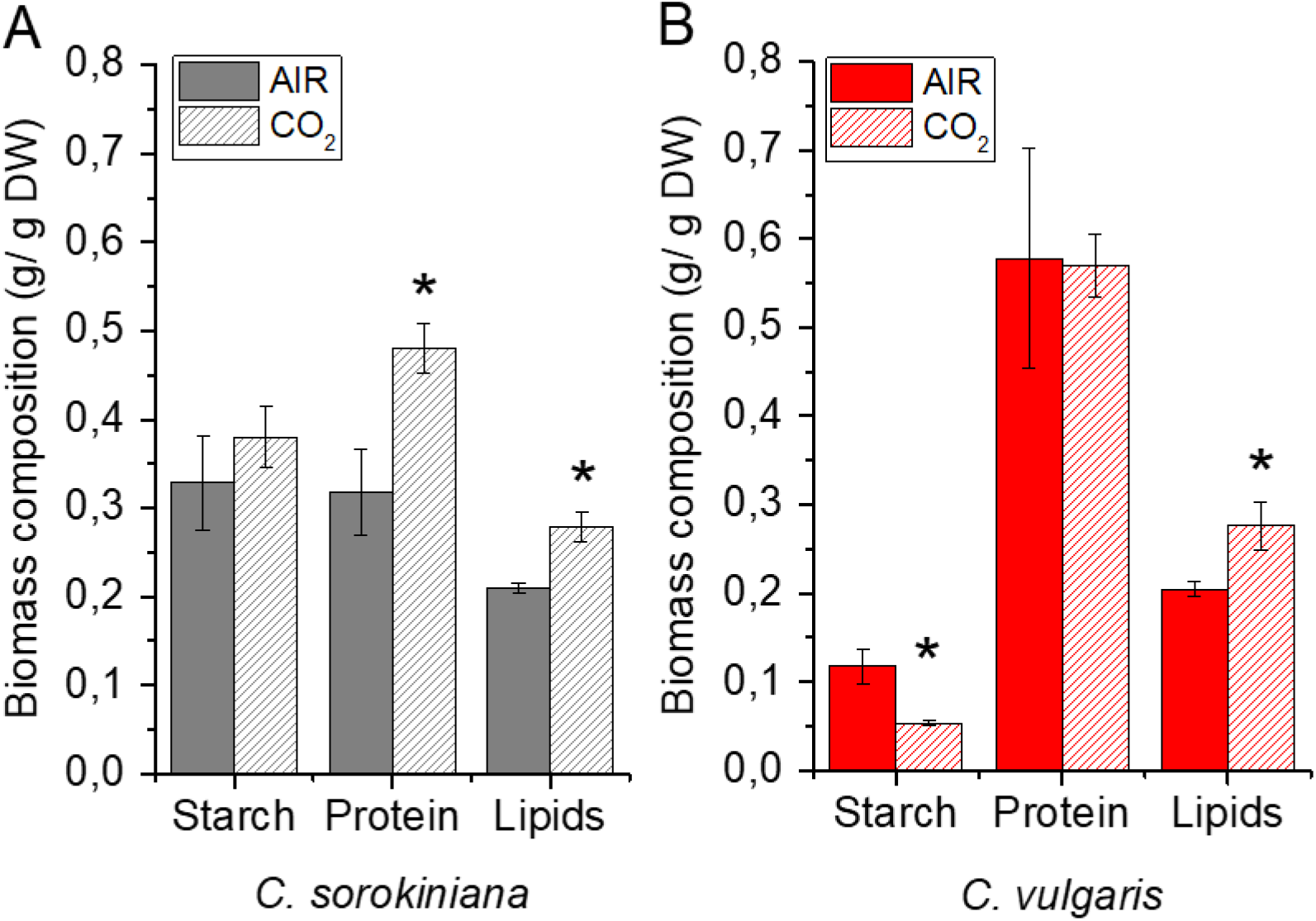
Starch, lipids and protein content in AIR *vs*. CO_2_. Relative starch, protein and lipid content per dry weight in *C. sorokiniana* (Panel A) and *C. vulgaris* (Panel B) in AIR *vs*. CO_2_ condition. Data are means of three biological replicates with standard deviation shown. Significantly different values in CO_2_ versus AIR are indicated by * (P < 0.05, n=3).

Fatty acid compositional and lipid class analyses revealed genus-specific response to varying CO_2_ availability. An increase in galactolipids under CO_2_ -replete conditions was observed in *C. sorokiniana* while triacylglycerol (TAG) was only increased in *C. vulgaris*. Differently, a decrease in phospholipids in CO_2_ could be noted in both genera (Figure 3). Specifically, a strong increase in monogalactosyldiacylglycerol (MGDG) and digalactosyldiacylglycerol (DGDG) per dry weight were observed in *C. sorokiniana*, when the availability of CO_2_ was high (CO_2_ condition). The decrease in galactolipids, major constituents of the thylakoid membranes in the chloroplast, suggests possible rearrangements at the level of the chloroplast organization. TAGs can derive from the recycling of pre-existing membrane glycerolipids as well as from *de novo* biosynthesis of fatty acids (Simionato et al., 2013). In *C. vulgaris*, all polar lipids were decreased in CO_2_ -replete conditions, likely due to a redirection of the metabolism to TAG biosynthesis. Thus, at high CO_2_ concentration *C. vulgaris* redirects its carbon flow from the storage of starch to the storage of TAGs, a more energy-dense carbon sink, while *C. sorokiniana* increased the fraction of lipids involved in thylakoid assembly. Interestingly, high CO_2_ availability led to an increase in the betaine lipid diacylglycerol *N,N,N*-trimethylhomoserine (DGTS) with a decrease in phosphatidylcholine (PC) in both species (Figure 3C-3F). The fatty acid profile of *C. vulgaris* and *C. sorokiniana* grown in AIR and CO_2_ is reported in Figure 3B-3E: *C. vulgaris* cells grown in high CO_2_ were characterized by a strong increase in palmitic acid (16:0), hexadecadienoic acid (16:2), stearic acid (18:0) and oleic acid (18:1) together with a decrease in 3-hexadecenoic acid i.e. 16:1 (3t). In *C. sorokiniana* cells grown in high CO_2_, an increase in palmitoleic acid (16:1 (9)), hexadecadienoic acid (16:2), hexadecatrienoic acid (16:3), linoleic acid (18:2) and α-linolenic acid (18:3) and a decrease in oleic acid (18:1) were observed. The strong increase in palmitic and oleic acid observed in *C. vulgaris* is consistent with the increased TAG accumulation observed in this species, being C16:0 and C18:1 fatty acids the main constituent of TAG in lipid droplets in microalgae (Siaut et al., 2011).

**Figure 3.**
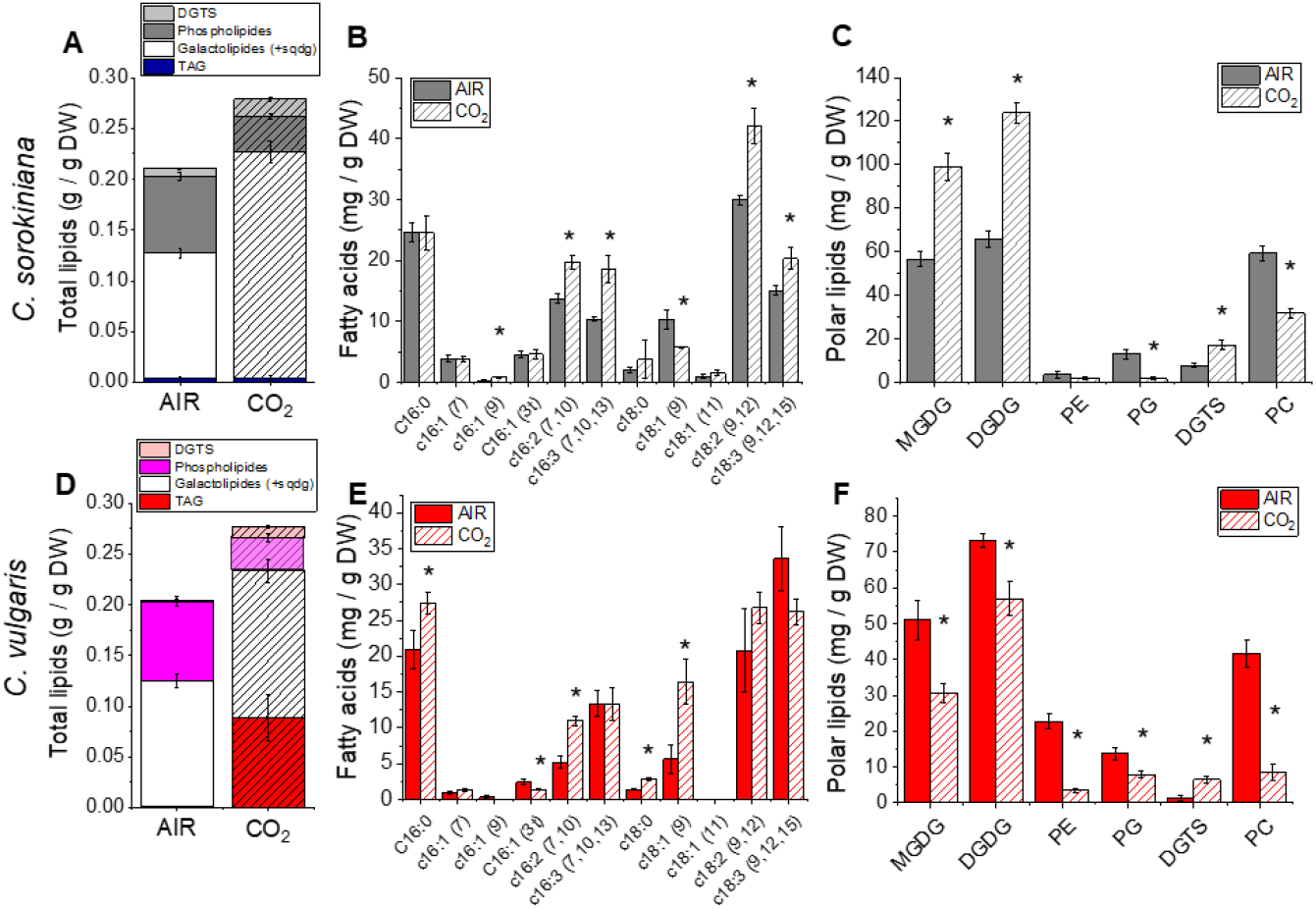
Lipid composition and fatty acids profile in AIR *vs*. CO_2_ conditions. Lipids composition of *C. sorokiniana* (panel A-C) and *C. vulgaris* (panel D-F) cells grown in AIR or CO_2_ conditions. Panel A and D: lipid composition in AIR *vs*. CO_2_ condition in terms of phospholipids, galactolipids, DGTS and triacylglycerol (TAG). Panel B and E: Fatty acids profile obtained by gas chromatography. Panel C and F: Polar lipid profile obtained by thin layer chromatography. Data are means of three biological replicates with standard deviation shown. Significantly different values in CO_2_ versus AIR are indicated by * (P < 0.05). MGDG, monogalactosyldiacylglycerol; DGDG, digalactosyldiacylglycerol; PG, phosphatidylglycerol; PE, phosphatidylethanolamine; PC, phosphatidylcholine; DGTS, diacylglycerol *N,N,N*-trimethylhomoserine.

### Photosynthetic properties of *Chlorella vulgaris* and *Chlorella sorokiniana* are differently influenced by CO_2_ availability

Photosynthetic properties of *Chlorella* species were investigated to determine the influence of different CO_2_ concentration on chloroplast metabolism. The amount of RUBISCO, being the key enzyme responsible for CO_2_ fixation in organic molecules, was first quantified. RUBISCO amount was similar in both species in the two conditions analysed on a cell basis (Figure S1), suggesting that RUBISCO accumulation was not tuned by CO_2_ concentration in *Chlorella*. This result is consistent with previous findings in soybean leaves exposed to different CO_2_ concentrations (Campbell et al., 1988). We then analysed PSII maximum quantum yield by measuring the fluorescence parameter F_v_ /F_m_, which is often used as a general indicator of the fitness of the culture. As reported in Figure 4C similar F_v_ /F_m_ values were found in both conditions, suggesting a minor, if any, impact of CO_2_ concentration in maximum PSII quantum yield. Interestingly, a strong decrease in chlorophyll (Chl) per cell was observed in *C. sorokiniana* grown in CO_2_ condition (Figure 4A). Moreover, in *C. sorokiniana* an increased Chl *a/b* ratio was evident in CO_2_ condition (Figure 4B): Chl *b* is bound only to the Light Harvesting Complex (LHC) subunits, the external antenna proteins of photosystems, while Chl *a* is bound to both antennae and core complex. A variation of the Chl *a*/*b* ratio suggests a change in the antenna/core complex stoichiometry, suggesting a rearrangement of the photosynthetic machinery. This adaptation in *C. sorokiniana* is consistent with results previously reported for the model alga *C. reinhardtii*, where a similar strong reduction of Chl content per cell was observed at high CO_2_ concentration (Polukhina et al., 2016). In contrast, this was not the case for *C. vulgaris*, where Chl/cell content was not significantly different in CO_2_ compared to AIR condition (Figure 4A).

**Figure 4.**
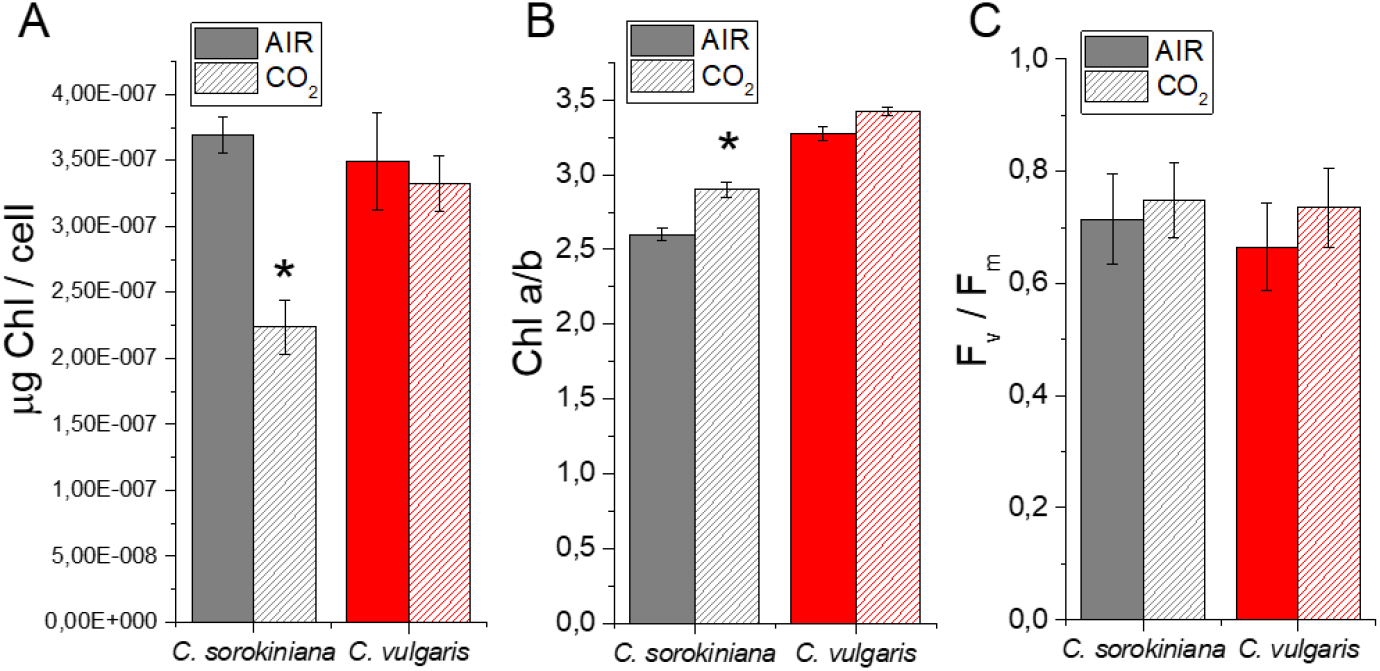
Chlorophyll content and PSII maximum yield in AIR *vs*. CO_2_ conditions. (A) Chlorophyll content per cell, (B) chlorophyll *a*/*b* ratio and (C) PSII maximum quantum yield expressed as F_v_ /F_m_ = (F_m_ -F_0_)/F_m_ in *C. sorokiniana* (grey colour) and *C. vulgaris* (red colour) in AIR (full colour) or CO_2_ (dash colour) condition. Data are means of three biological replicates with standard deviation shown. Significantly different values in CO_2_ versus AIR are indicated by * (P < 0.05).

To further investigate remodelling of the components of photosynthetic complexes, PSI/PSII ratio and LHCII/PSII ratio were evaluated by immunoblot analysis (Figure 5A and 5B). In *C. sorokiniana* both PSI/PSII and LHCII/PSII ratio were decreased in CO_2_ compared to AIR condition, while no significant differences were observed in *C. vulgaris* (Figure 5C and 5D). PSI photochemical activity can be measured by transient absorption: oxidation of the reaction centre of PSI, called P700, causes an increased absorption at 830 nm, thus following transient absorption kinetics at this wavelength it is possible to evaluate the PSI activity. Transient absorption measurements at 830 nm were performed in *Chlorella* species in the presence of DCMU, which inhibits linear electron transport from PSII to PSI, and ascorbate and methyl viologen as an electron donor and acceptor, respectively. In the case of *C. sorokiniana* a ∼40% reduction of the maximum PSI activity was observed in CO_2_ condition compared to the AIR case (Figure 5E), consistent with the reduced PSI/PSII ratio observed by immunoblot analysis (Figure 5D).

**Figure 5.**
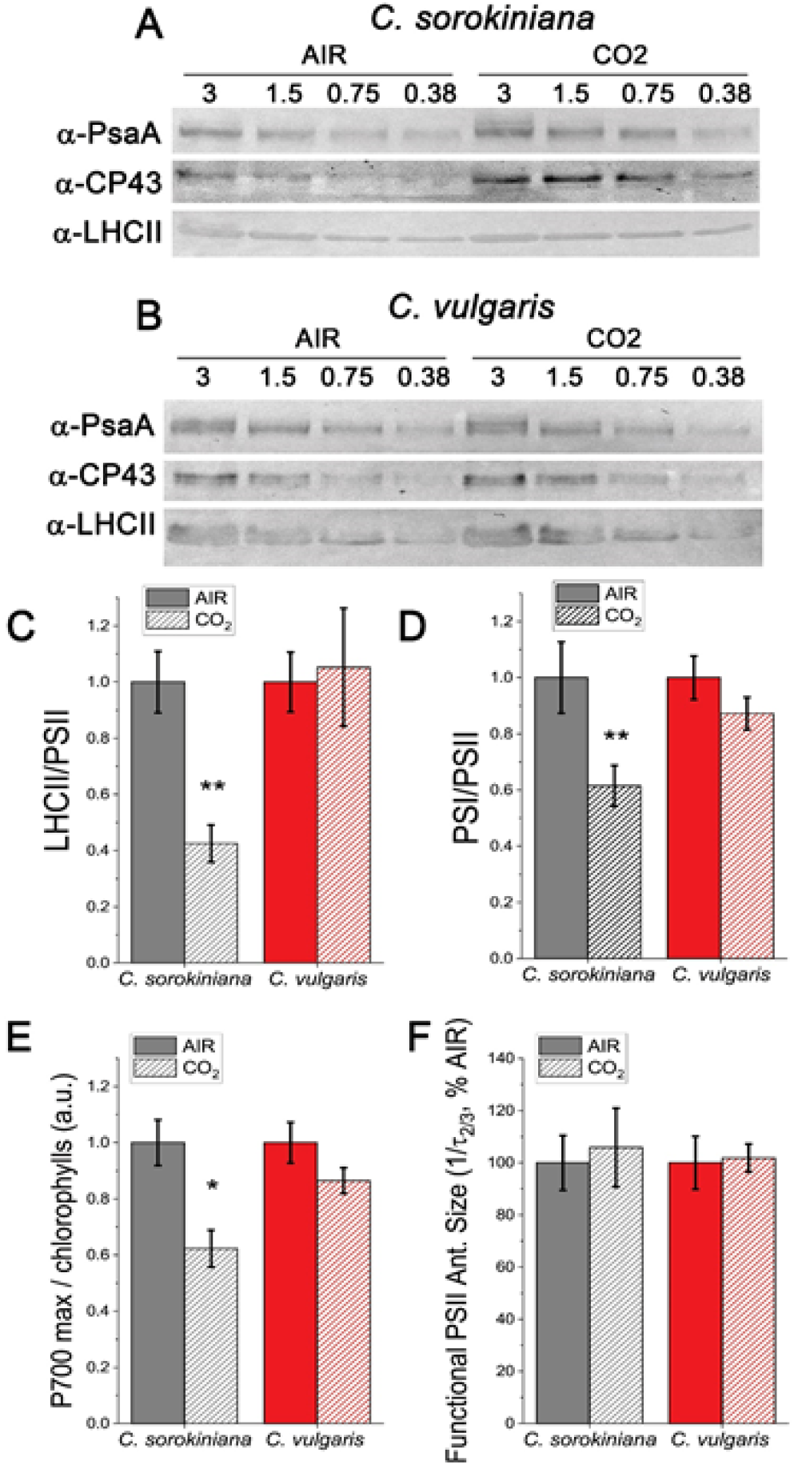
Analysis of PSI, PSII and LHCII content by immunoblots, P700 activity and functional PSII antenna size. (A, B) Immunoblot analysis of PSI (α-PsaA antibody), PSII (α-CP43 antibody) and LHCII (α-LHCII antibody). Loading was performed on a chlorophyll basis: total µg chlorophylls loaded in each lane is reported on the top of Panel A and B. (C, D) PSI/PSII (C) and LHCII/PSII (D) ratios calculated by densitometry of immunoblot signals for *C. sorokiniana* (panel A, grey colour) and *C. vulgaris* (panel B, red colour) in AIR (full colour) or CO_2_ (dash colour) condition. (E) Maximal P700 oxidation on a chlorophyll basis in *C. sorokiniana* (left, grey colour) and *C. vulgaris* (right, red colour) in AIR (full colour) or CO_2_ (dash colour) normalized to AIR condition. (F) Functional antenna size of the photosystem II (1/τ_2/3_) normalized to AIR condition in *C. sorokiniana* (grey colour) and *C. vulgaris* (red colour). Data are means of three biological replicates with standard deviation shown. Significant different values in CO_2_ versus AIR are indicated by ** (P < 0.01) and by * (P < 0.05).

The reduced LHCII/PSII ratio observed in *C. sorokiniana* is consistent with the increased Chl *a*/*b* ratio (Figure 4B). To investigate whether the different LHCII/PSII ratio induced by CO_2_ availability in *C. sorokiniana* influences a functional light harvesting capacity of PSII, as previously observed in *C. reinhardtii* (Berger et al., 2014), fast Chl *a* fluorescence emission spectrum in presence of DCMU was measured for both *C. vulgaris* and *C. sorokiniana* grown in AIR and CO_2_ condition. Light harvesting capacity, or functional antenna size, of PSII was indeed reported to be inversely proportional to the time required to reach 2/3 of the maximum Chl *a* fluorescence emission upon inhibition of PSII electron transport activity (Malkin et al., 1981). As shown in Figure 5F and Figure S2 no differences in PSII functional antenna size were observed in *C. sorokiniana* or in *C. vulgaris* depending on CO_2_ availability. This result indicates that the reduction of LHCII/PSII ratio measured in *C. sorokiniana* did not affect the PSII light harvesting capacity, being thus likely related to LHCII subunits poorly connected to PSII.

LHC complexes are also involved in the process called state transitions, where a fraction of the antenna complexes bound to PSII moves to PSI to maintain the excitation balance between the two photosystems. This process is triggered in *C. reinhardtii* by LHC phosphorylation catalysed by a kinase enzyme called STT7 (Depege et al., 2003). State 1 (S1) or State (S2), being respectively the conditions with minimum or maximum migration of LHCII to PSI, can be induced by consuming or increasing the reducing power in the chloroplast as described in (Fleischmann and Rochaix, 1999). In particular, S1 or S2 state can be measured on whole cells by measuring Chl fluorescence emission at 77K, where both PSI and PSII emission are detectable. As reported in Figure 6 both *C. vulgaris* and *C. sorokiniana* were able to undergo state transitions in both AIR and CO_2_ with an increased PSI contribution in S2 compared to S1. However, *C. sorokiniana* cells grown in CO_2_ exhibited an increased capacity for state transitions compared to cells grown in AIR. When 77K fluorescence emission in S1 or S2 were compared to 77K fluorescence of cells directly harvested in their growing conditions, it was possible to observe that both *C. vulgaris* and *C. sorokiniana* were essentially growing in S2 state in AIR condition. A different behaviour was instead observed in CO_2_ condition between the two species herein analysed: while *C. vulgaris* cells were characterized by an intermediate state between S1 and S2, *C. sorokiniana* was essentially in S1. These results indicate that the reduction in *C. sorokiniana* grown in CO_2_ of LHCII/PSII ratio was related to a decrease in LHCII preferentially connected in AIR to PSI, suggesting a compensatory regulation between antenna regulation and state transitions.

**Figure 6.**
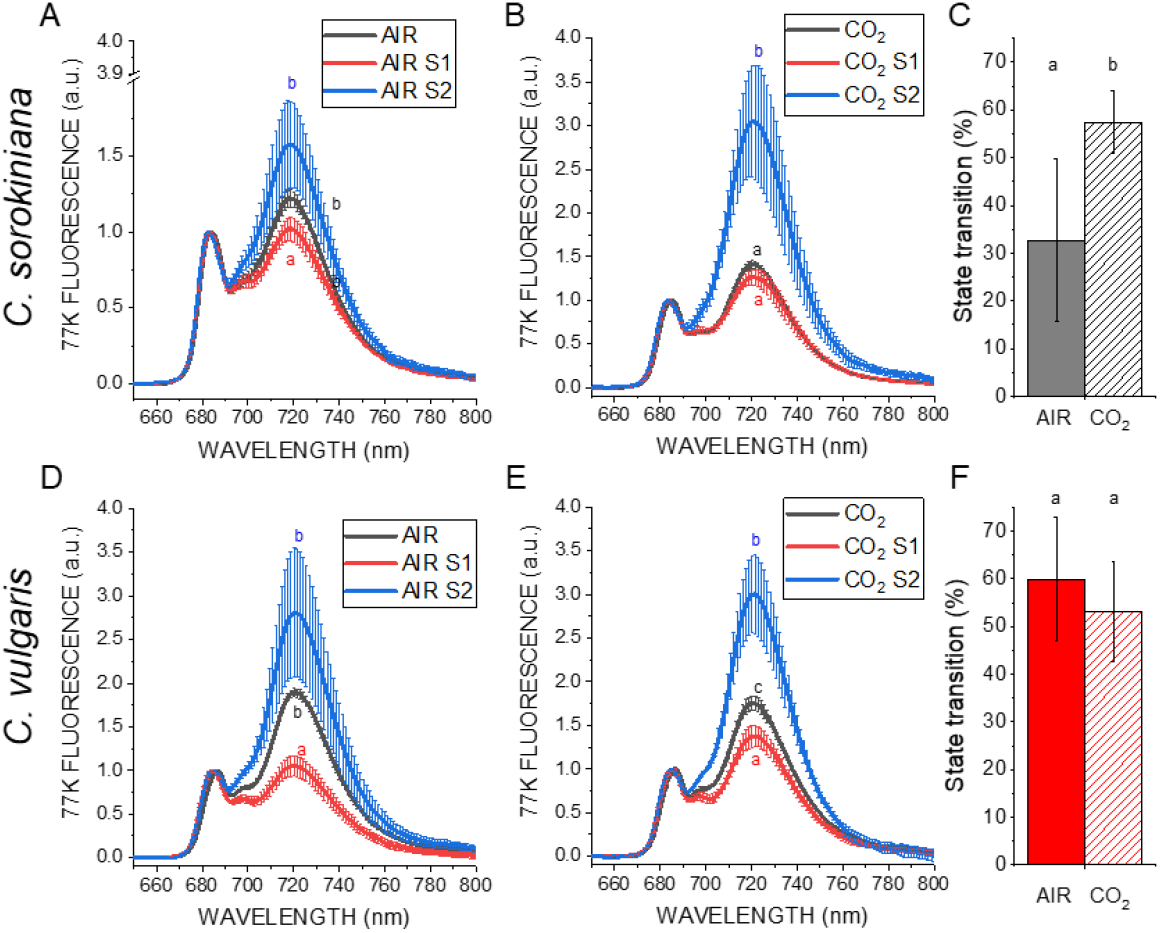
State transitions in AIR *vs*. CO_2_. State transition analysis by 77K fluorescence emission spectra in state 1 (S1, red lines) or state 2 (S2, blue lines) conditions in *C. sorokiniana* (A-C) and *C. vulgaris* (D-F) in AIR (A, D) or CO_2_ (B, E). S1 was induced by shaking vigorously cells in a low light (□5μmol m^2^ s^-1^) with 10µm of DCMU for at least 15 min to oxidize the plastoquinone pool while S2 was induced by adding 250 μm sodium azide to inhibit mitochondrial respiration and to reduce the plastoquinone pool as described in Fleischmann et al. 1999. Black lines are related to cells harvested in the different growing conditions (AIR or CO_2_) and dark adapted for 1 minute before freezing at 77K. Maximum capacities for state transitions were then quantified from the maximum fluorescence emission at 720 nm as (F_S2_ -F_S1_)/F_S2_. Data reported are means of three biological replicates with standard deviation shown. Significant difference in CO_2_ *vs*. AIR are indicated with different letters (a, b, or c, P < 0.05).

Photosynthetic electron transport is coupled to the formation of a proton gradient across the thylakoid membrane, exploited by ATPase as proton motif force to produce ATP (Tikhonov, 2013). ATPase content on Chl basis and proton-motive force (*pmf*) upon exposure to different light intensity were evaluated in CO_2_ and AIR grown cells (Figure 7). The *pmf* can be estimated by measuring the light dependent-electrochromic shift of carotenoid absorption (Bailleul et al., 2010). In this case the behavior of the two *Chlorella* species was similar with a reduced *pmf* in CO_2_ compared to AIR condition. At the same time, an increase in ATPase content under CO_2_ condition was detected for both *C. vulgaris* and *C. sorokiniana*. Likely the higher level of ATPase in CO_2_ condition improved proton movement back to the stroma resulting in reduced *pmf* and higher ATP production in CO_2_ condition. Furthermore, we investigated the influence of cyclic electron flow (CEF) around PSI measuring electrochromic shift (ECS) in the presence of DCMU inhibiting PSII and thus linear electron flow. Only a 2-7% of residual *pmf* was detected in DCMU treated samples, indicating a low level of CEF operating in *C. vulgaris* and *C. sorokiniana*, not significantly influenced by CO_2_ concentration.

**Figure 7.**
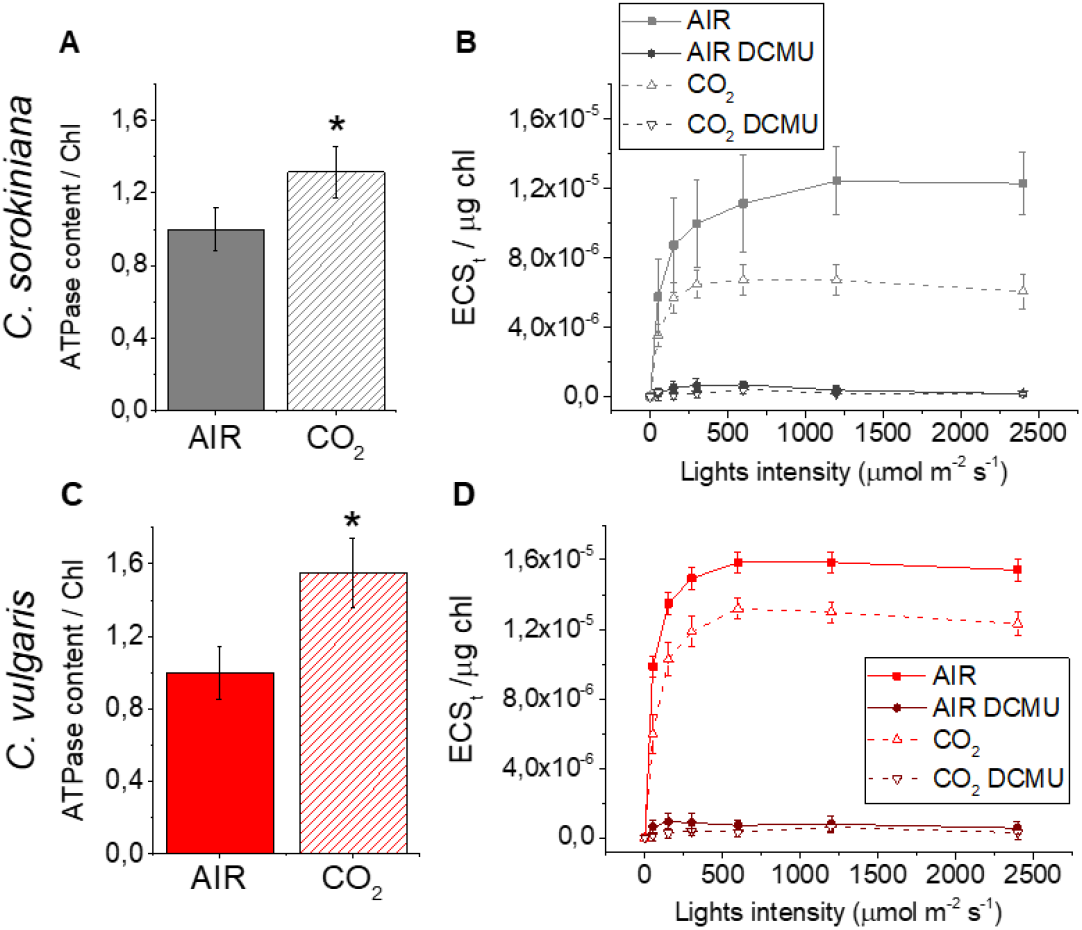
ATPase content and electrochromic shift in AIR *vs*. CO_2_. Immunoblot analysis of ATPase content (atpC subunit antibodies) and ECS measurements in *C. sorokiniana* and *C. vulgaris* in AIR (solid line,) or CO_2_ (dashed line) condition. ECS results in presence of DCMU are also reported with open symbol. Data are means of three biological replicates with standard deviation shown. Significantly different values in CO_2_ versus AIR are indicated by * (P < 0.05).

PSI is a plastocyanin-ferredoxin oxidoreductase that reduces NADP^+^ to NADPH by a ferredoxin–NADP^+^ reductase (FNR) enzyme. In parallel, the mitochondrial respiratory electron transport chain oxidase NADH releasing NAD^+^. Chloroplasts and mitochondria communicate to balance the NAD(P)^+^/NAD(P)H pool (Johnson and Alric, 2013; Dang et al., 2014; Uhmeyer et al., 2017). We evaluated the light dependent NADPH formation rate by following NAD(P)H fluorescence changes upon exposure to actinic light of 300 μmol photons m^−2^ s^−1^ for 120s. It is important to note that NADH and NADPH cannot be distinguished by fluorescence, both contributing to the signals herein detected. In both species either in AIR or CO_2_ conditions the rates of NAD(P)H fluorescence during actinic light exposure were negative, indicating that NAD(P)H consumption exceeds light dependent NADPH production (Figure 8). It’s interesting to observe that in *C. sorokiniana* the same balance between NAD(P)H formation and consumption was maintained comparing AIR *vs*. CO_2_ condition, while in *C. vulgaris* a higher rate of NADPH consumption was observed in CO_2_ condition.

**Figure 8.**
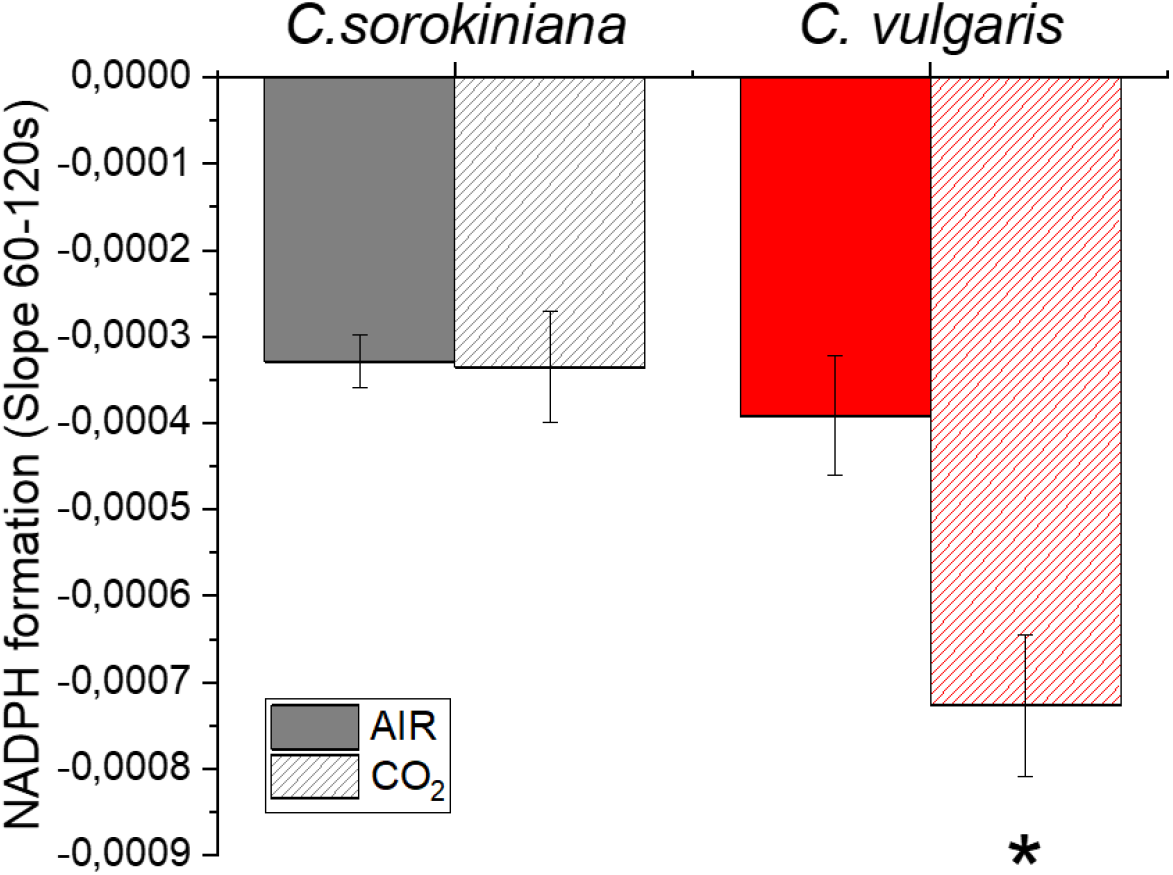
NAD(P)H formation rate in AIR *vs*. CO_2_. Light dependent rate of NAD(P)H formation upon exposure to light (300 μmol photons m^−2^ s^−1^) for 120s in *C. sorokiniana* (grey colour) and *C. vulgaris* (red colour) in AIR (full colour) or CO_2_ (dash colour) condition. The data reported were calculated from the slope of the NAD(P)H fluorescence emission curve upon exposure to actinic light. Data are means of three biological replicates with standard deviation shown. Significantly different values in CO_2_ versus AIR are indicated by * (P<0.05).

### The response of mitochondrial respiratory pathways to CO_2_ availability

Mitochondrial respiration is a fundamental process that allows producing ATP while releasing NAD^+^ that can return to the chloroplast. The mitochondrial electron transport chain, also called cytochrome pathway, includes an ATP synthase complex, called also complex V, and four oxidoreductase complexes that oxidise the reducing power and produce ATP thank to the electrochemical gradient that is formed across the membrane. In addition, an alternative oxidase (AOX) might operate directly coupling ubiquinol oxidation with the reduction of O_2_ to H_2_ O serving as an alternative route bypassing the electron transport chain thus dramatically reducing the energy (ATP) yield. AOX was reported having a role in the protection mechanism for the respiratory chain (Boekema and Braun, 2007; Vanlerberghe, 2013).

The contribution of cytochrome and alternative pathways (Figure 9) was investigated by measuring the dark respiration in the presence of two specific inhibitor: SHAM (salicylhydroxamic acid) that inhibits AOX and so the alternative pathway, and myxothiazol that locks the complex III therefore blocking the cytochrome pathway (Dang et al., 2014). We observed that the total dark respiration on a cell basis is essentially unaffected in *C. sorokiniana*, while a strong reduction was reported for *C. vulgaris* in CO_2_ condition. In both species a reduction of the fraction of dark respiration operating through AOX was evident, leading to an increased efficiency of ATP production by NADH oxidation through the cytochrome pathway.

**Figure 9.**
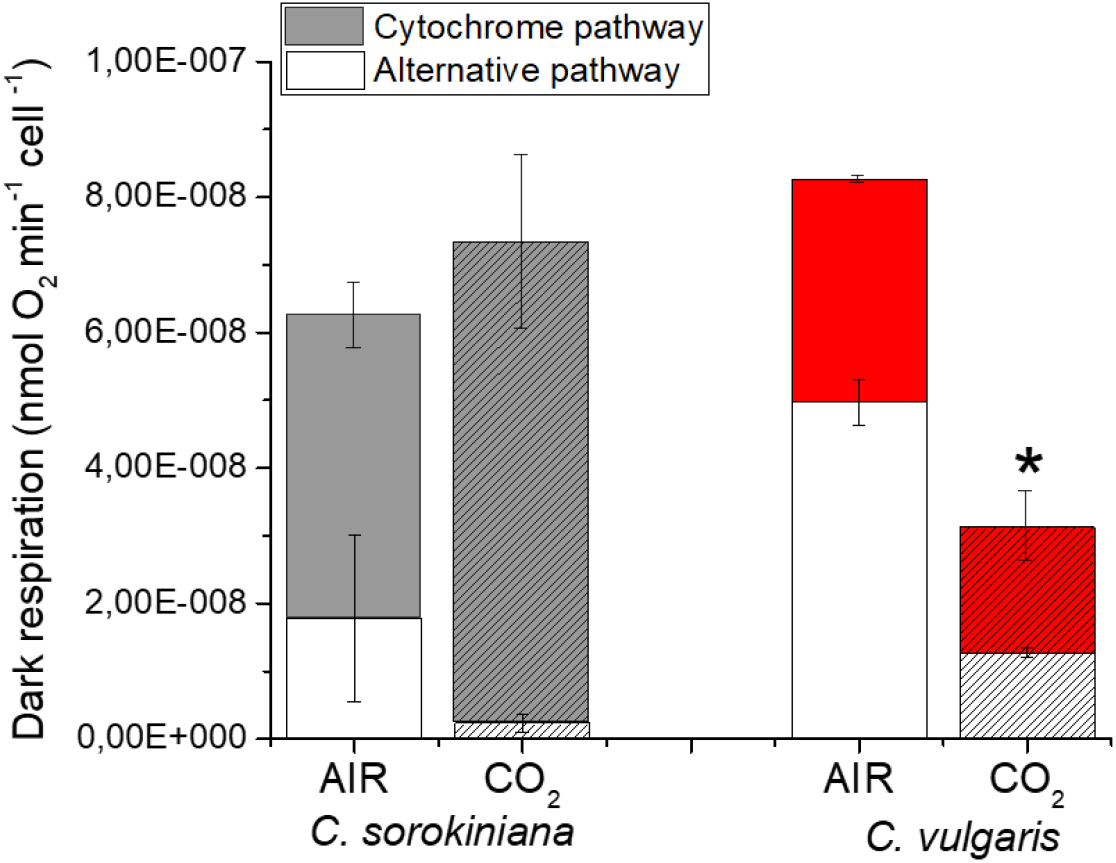
Dark respiration in *C. sorokiniana* and *C. vulgaris* in AIR *vs*. CO_2_ conditions. The relative contribution of cytochrome (filled bars) and alternative respiration (empty bars) was reported normalized to cell content. Data are means of three biological replicates with standard deviation shown. Significant difference in CO_2_ versus AIR are indicated by * (P < 0.05).

### NAB1-like proteins in *C. vulgaris* and *C. sorokiniana*

Acclimation to different carbon availability has been reported in *C. reinhardtii* to involve the translational repressor NAB1. NAB1 acts as a molecular switch triggered by the redox state of the cell, which is in turn strongly influenced by the carbon availability, repressing the translation of specific transcripts, including those for LHC subunits. To evaluate the possible conservation of NAB1 in the *C. vulgaris* and *C. sorokiniana* species, BLAST search was performed using the *C. reinhardtii* protein sequence as query. It is important to note that functional NAB1 in *C. reinhardtii* is composed of a Cold-shock domain (CSD) at the N-terminus and a RNA recognition motif (RRM) at the C-terminus (Mussgnug et al., 2005). Among the putative protein sequences identified by BLAST, only in *C. vulgaris* the g211.t1 locus containing both CSD and RRM domains were conserved. Both CSD and RRM domain were found also in a predicted mega-protein in *C. sorokiniana* (CSI2_123000002385-RA) where two additional V-ATPase proteolipid subunit C-like domains were present, suggesting that this polypeptide has likely different functions compared to the *C. reinhardtii* NAB1 (Figure S3). Consistent with the bioinformatic analysis, immunoblot analysis using anti-NAB1 antibody revealed a band at the expected molecular weight (27 kDa) in both *C. reinhardtii* and *C. vulgaris* but not in *C. sorokiniana* (Figure S4). However, while the accumulation of NAB1 was increased in AIR compared to CO_2_ in *C. reinhardtii* condition, as previously described, a similar level of NAB1-like protein was observed in *C. vulgaris* cells grown in AIR or CO_2_ (Figure S4). As recently reported, the transcriptional upregulation of NAB1 in *C. reinhardtii* at low CO_2_ availability is related to the transcription factor LCRF which belongs to the Squamosa promoter binding protein (SBP) family of transcription factors (Blifernez-Klassen et al., 2021). Possible homologous of *C. reinhardtii* LCRF was thus searched in *C. vulgaris* and *C. sorokiniana* by BLAST search. Consistently with the absence of a NAB1-dependent adaptation mechanisms, no homologous of *C. reinhardtii* LCRF could be found in in *C. vulgaris* and *C. sorokiniana*.

Sequence analysis of the identified NAB1-like protein in *C. vulgaris* demonstrated that among the key residues involved in NAB1 activity regulation in *C. reinhardtii* (Wobbe et al., 2009; Blifernez et al., 2011; Berger et al., 2016), only Cys181 and Arg92 were conserved, while Cys226 or Arg90 are substituted by a valine and a serine residue, respectively, in the NAB1-like subunit in *C. vulgaris* (Figure S5). It is important to note that the substitution of Cys226 in *C. reinhardtii* NAB1 arrested the protein in its active state and abolished the redox control mechanism leading to a pale green phenotype in strains exclusively expressing NAB1_Cys226Ser_ (Wobbe et al., 2009). As reported in Figure S2, when comparing the functional antenna size of *C. reinhardtii, C. vulgaris* and *C. sorokiniana* grown in AIR or CO_2_ conditions, a smaller light harvesting capacity was evident in the *Chlorella* species compared to *C. reinhardtii* in CO_2_, while similar PSII antenna size was observed in AIR. This result suggests that NAB1 dependent adaptation of PSII antenna size to different CO_2_ concentration is not conserved in *C. vulgaris* and *C. sorokiniana*, because either the protein is not expressed (*C. sorokiniana*) or the redox control is not functional (*C. vulgaris*). As another important difference, the NAB1 homolog in *C. vulgaris* does not accumulate upon CO_2_ limitation.

## DISCUSSION

Atmospheric CO_2_ concentration has significantly increased over the last 100 years and is continuing rising at an unprecedented speed. This greenhouse gas strongly contributes to climate change and global warming leading to a potential severe environmental crisis. Microalgae are promising platforms to capture CO_2_, possibly integrating microalgae cultivation with CO_2_ recovery from flue gasses, thus reducing industry derived CO_2_ emission and carbon footprint. For this reason, understanding the cell acclimation process involved in CO_2_ metabolism is crucial to develop new strategies for improving the ability of microalgae to acquire and accumulate carbon. In this work we focused on two of the most promising species for microalgae cultivation at industrial level, *C. sorokiniana* and *C. vulgaris* (Li et al., 2013; Sun et al., 2016; Bernaerts et al., 2019; Camacho et al., 2019; Niccolai et al., 2019). They were grown in airlift photobioreactors under atmospheric level of CO_2_ (□0.04% CO_2_, AIR condition) or 3% CO_2_ (CO_2_ condition). We discuss the metabolic consequence of the different photosynthetic and respiratory responses to high or low CO_2_ levels in two *Chlorella* genus.

### CO_2_ availability boosts biomass accumulation

Increasing CO_2_ supply boosted (□260% increase) in biomass yield in both species. Interestingly, differential response was observed between the two strains in terms of biomass composition (i.e. protein, starch and lipid amount). In *C. vulgaris* a decrease in starch accumulation and an increase in lipid accumulation, in particular TAG, were detected. This suggests a redirection of the energy reserves from starch to TAG accumulation, a class of macromolecules with a higher energy content per gram, indicating an improved light energy conversion. In *C. sorokiniana* not only lipids, but also protein content increased, the latter being an additional carbon sink in cells grown at high CO_2_ concentration.

### Lipid remodelling and CO_2_ availability

Both *Chlorella* genus showed an increase in total lipid amount under high CO_2_, and they further remodelled their fatty acid as well as lipid class composition albeit in different ways. Phospholipids were reduced whereas the betaine lipid DGTS were increased in both species by high CO_2_ level. Betaine lipid, a non-phospholipid, has been observed often increased during phosphate limitation, presumably replacing the function of phospholipids in cell membranes (Riekhof et al., 2014; Murakami et al., 2018; Hidayati et al., 2019). For still unknown reasons, a similar response between P limitation and high CO_2_ was observed. We can speculate that increased CO_2_ availability required phosphate reallocation to the different macromolecules produced, requiring partial phospholipids substitution with betaine lipid DGTS.

In *C. vulgaris*, the increase in lipid content was mostly due to an increase in TAG, whereas, in *C. sorokiniana* the increase in total lipids was mainly due to an increase in the two galactolipids (MGDG and DGDG), the major lipids of photosynthetic membranes (Li-Beisson et al., 2019). The differential response in lipid classes in the two strains to high CO_2_ level is further supported by fatty acid compositional alterations. Among other reasons, lipid compositional changes are mostly likely results of altered redox status brought about by differential chloroplast and mitochondrial energetic activities in response to varying CO_2_ availability in the two *Chlorella* genus (Figure 6, Figure 8) (Burlacot et al., 2019).

### Thylakoid reorganization and photosynthetic activity under varying CO_2_

The increase in DGDG and MGDG, surprisingly, was observed together with a reduction in Chl content per cell in *C. sorokiniana* under high CO_2_. Nevertheless, the observed reduction of Chl content per cell in *C. sorokiniana* grown in CO_2_ condition compared to AIR was in line with the results obtained previously in *C. reinhardtii* (Polukhina et al., 2016), while this was not the case for *C. vulgaris*, where Chl content is independent from CO_2_ availability. Again, similar to *C. reinhardtii, C. sorokiniana* grown in high CO_2_ was characterized by reduction of LHCII/PSII content and a reduction in PSI/PSII ratio, while these adaptations were not observed in *C. vulgaris*. The reduced LHCII/PSII content observed in *C. sorokiniana* grown in CO_2_ condition did not affect the functional antenna size of PSII: different from previous observation in *C. reinhardtii*, functional antenna size of PSII was not influenced by CO_2_ availability in both *Chlorella* species herein investigated (Figure 5F, Figure S2). In *C. reinhardtii* it was indeed reported that high CO_2_ availability caused an increase in the functional antenna size of PSII, due to suppressed accumulation at high CO_2_ concentration of NAB1, the translation repressor specific for LHCII encoding mRNAs (Berger et al., 2014). Here, we report the identification of a NAB1-like protein in *C. vulgaris* only, which however is not differently expressed in AIR *vs*. CO_2_ conditions. Accordingly, homologous protein of the transcription factor LCRF, the transcriptional regulator of NAB1 recently identified in *C. reinhardtii* (Blifernez-Klassen et al., 2021) could not be found in either *C. vulgaris* or *C. sorokiniana*. Moreover, in the *C. vulgaris* NAB1-like subunit only Cys181 and Arg92 residues are conserved among the two cysteine (Cys181, Cys226) and two arginine (Arg90, Arg92) residues reported in *C. reinhardtii* NAB1 to be involved in its activity as translational repressor. Taken together, the NAB1-dependent regulation of PSII antenna size at different CO_2_ concentration in *C. reinhardtii* is absent in both *C. sorokiniana* and *C. vulgaris*, where NAB1 homolog was respectively not identified or is missing the crucial redox control of its translational inhibition activity (Figure S3, Figure S4). Indeed, the PSII antenna size of both *C. sorokiniana* and *C. vulgaris* was not affected by different CO_2_ availability, being similar to the PSII antenna size measured in the case of *C. reinhardtii* grown in AIR (Figure S2). Regulation of PSII antenna size by NAB1 is thus an acclimation mechanism finely controlled by the redox state of the cell which is not conserved among *Chlorophyta*.

It is important to note that the effect of CO_2_ availability on *C. reinhardtii* LHCII/PSII ratio is still under debate, with Polukhina and coworker reporting a general reduction of LHCII/PSII content in *C. reinhardtii* grown in high CO_2_, in parallel with a decrease in PSI/PSII ratio (Polukhina et al., 2016). Considering the possibility of LHCII proteins to function as PSI antenna, it was not excluded by Polukhina and coworkers that the decrease of LHCII content per PSII observed at high CO_2_ might be mainly related to the amount of LHCII proteins acting as PSI antenna. Here we report a similar acclimation mechanism only in the case of *C. sorokiniana*, with the difference that PSII antenna size was not modulated by CO_2_ availability.

### Cellular redox balance and CO_2_ availability

Cellular reducing power is crucial for the overall carbon flow and cell metabolism: catabolic process generate reducing power which can be used by oxidative phsophorylation to generate ATP or by the anabolic pathways. In photosynthetic organisms, the NADP^+^/NADPH balance influence both the light phase of photosynthesis and the carbon fixation reactions. In general it is possible to hypothesize that the increased capacity of Calvin Benson cycle to regenerate NADP^+^ and ADP, thanks to the increased CO_2_ availiability, trigger the light phase of photosynthesis in order to keep the NADPH/NADP^+^ ratio similar to the AIR condition, as reported in Figure 8: this occurs by increasing the total amount of PSII compared to PSI and relatively redistributing the excitation pressure among PSI and PSII reducing the excitation pressure at the level of the former reducing the LHCII content bound to PSI. This acclimation process would explain the reduced LHCII/PSII content despite the similar PSII antenna size observed. The absence of such acclimation mechanisms in the case of *C. vulgaris* could be at the base of the strong reduction of NAD(P)H/NAD(P)^+^ ratio observed at high CO_2_ concentration, as a consequence of increased NADPH consumption by the Calvin Benson cycle. Alteration of RUBISCO content was not detected (Figure S1), suggesting an enhanced RUBISCO activity due to the higher availability of substrate rather than an upregulation of enzyme to exploit the higher CO_2_ availability. CO_2_ fixation requires both ATP and NADPH: in both *Chlorella* species a decrease of *pmf* and an increased ATPase content was measured in CO_2_ condition (Figure 6). Likely, the higher level of ATPase in CO_2_ condition prevents the accumulation of the electrochemical gradient, suggesting a higher ATP production.

Interestingly, dark respiration is differentially regulated in the two *Chlorella* species: in *C. sorokiniana* total dark respiration was similar in AIR compared to CO_2_ condition, with an increased NADH oxidation through the cytochrome pathway and reduced AOX activity. Accordingly, in *C. sorokiniana* we observed the same balance of the NAD(P)H redox state: the rearrangements of the photosynthetic machinery in CO_2_ condition improved the pool of NADPH and ATP, likely matching the increased substrate (CO_2_) availability for sugar production by the Calvin Benson cycle. In contrast, in *C. vulgaris* a strong reduction of dark respiration in CO_2_ condition was evident, despite an increase of cytochrome/alternative pathway ratio. Additionally, in *C. vulgaris* there was a higher NAD(P)H consumption in CO_2_ suggesting that chloroplast acts as a sink of reducing power subtracting them from the mitochondrion. Moreover, the relative reduction of starch accumulation and the increase of TAG suggested a redirection of photosynthates to other metabolic pathways. Consumption of triose phosphates by the glycolytic pathway leading the acetyl-CoA production could be a possible link between reduced starch accumulation and increased TAG content. Indeed, an increase of acetyl-CoA and reducing power was reported at the base of the increased TAG accumulation observed in the diatom *Phaeodactylum tricornutum* (Valenzuela et al., 2012; Yang et al., 2013; Li-Beisson et al., 2019).

## CONCLUSIONS

High CO_2_ availability caused an increased biomass accumulation in both *C. vulgaris* and *C. sorokiniana*, likely related to increased photosynthates production. The increased carbon assimilation in high CO_2_ redirected the metabolism toward biosynthesis of lipids (TAG) in *C. vulgaris*, and proteins in *C. sorokiniana*, respectively. Increased carbon fixation at high CO_2_ concentration requires an increased NADPH and ATP availability: while increased ATPase content and reduced *pmf* suggesting indeed an increased ATP regeneration in both *C. vulgaris* and *C. sorokiniana*, the increased NADPH requirement was differently satisfied in *C. vulgaris* and *C. sorokiniana* in CO_2_. In the case of *C. vulgaris*, chloroplast acted as a sink for reducing power, inducing consequently a reduced NADH availability for mitochondrial respiration, reducing in particular the relative contribution of alternative pathways, not related to ATP biosynthesis. In *C. sorokiniana* a reduced PSI/PSII ratio and reduced LHCII binding to PSI was observed in CO_2_, allowing an increased electron flow toward NADP^+^ to NADPH regeneration and ensuring a similar NAD(P)H/NAD(P)^+^ ratio in both AIR or CO_2_ conditions. A summary of the adaptation to CO_2_ condition is shown in Figure S6. Elucidation of the molecular rearrangements in enriched CO_2_ condition could be useful to develop strategies to improve in these species and in other microalgae of industrial interest their carbon assimilation efficiency to improve sustainability, biomass yield and productivity of specific compounds.

## MATERIAL AND METHODS

### Microalgae cultivation

*C. sorokiniana* UTEX 1230 and *C. vulgaris* 211/11P strain (Culture Collection of Algae at Goettingen University CCAP 211/11P strain) cells were grown in the Multi-Cultivator MC 1000 tubes aerated with air or with 3% CO_2_ -enriched air obtained by a gas mixing system. Cells were grown in BG11 medium starting from 1*10^6^ cell/ml at 300 μmol photons m^−2^ s^−1^ (Allen and Stanier, 1968). Cell number was determined Countless®II FL automated cell counter (Thermo Fisher). The cell density was automatically monitored every ten minutes by measuring the absorption at 720 nm. For physiological measurements, cultures were harvested during the exponential growth phase. At the end of the growth curve the dry weight determination was performed: cell culture was harvested by centrifugation at 4500g for 5 min at 20°C then drying in a lyophilizer for 48h and then net dry weight was calculated.

### Biomass composition analysis

Lipid, starch and protein content of the biomass harvested at the end of the exponential phase were analyzed as previously reported in Cecchin et al. 2019.

### Photosynthetic parameters and pigments extraction

The pigments were extracted with 100% DMSO at 60°C in dark conditions and measured with Jasco V-550 UV/VIS spectrophotometer. Proton motive force upon exposure to different light intensities was measured by Electrochromic shift (ECS) with MultispeQ v2.0 (PhotosynQ) according to Kuhlgert et al. 2016 and normalized to the Chl content of the sample. PSII activity was analyzed by fluorescence measurements on whole cells using a Dual-PAM 100 instrument (WALZ). 77K fluorescence emission spectra were acquired with a charge-coupled device spectrophotometer (JBeamBio) as previously described (Allorent et al., 2013). State transitions were measured on whole cells induced to state 1 or state 2 as described in Fleischmann et al. 1999. PSII functional antenna size was measured from fast Chl induction kinetics induced with a red light of 11 μmol photons m^-2^ s^-1^ on dark-adapted cells incubated with 50 μM DCMU (Malkin et al., 1981). The reciprocal of time corresponding to two-thirds of the fluorescence rise (τ_2/3_) was taken as a measure of the PSII functional antenna size (Malkin et al., 1981). P700 activity was measured with the DUAL-PAM-100 (Heinz-Walz) following the transient absorption at 830 nm upon exposure to actinic light. Maximum P700 activity was measured after a pulse of saturating light inwhole cells treated with DCMU (3-(3,4-dichlorophenyl)-1,1-dimethylurea), ascorbate, and methyl-viologen, as described in Bonente et al. 2012. The formation rate of NADPH was determined with the NADPH/9-AA module of the Dual-PAM 101 (Schreiber and Klughammer, 2009). Cells were harvested and resuspended in BG11 medium with 10% of Ficol to reduce cells precipitation. Measurement was performed as described in Schreiber and Klughammer 2009 at the same light intensity of growth (300 μmol photons m^−2^ s^−1^). The slope during the light phase, between 60-120s, was used to determine the rate of NADPH formation.

### SDS-PAGE and immunoblotting

SDS-PAGE and immunoblotting were performed as described in Bonente et al. 2011. The following antibodies obtained from Agrisera company (https://www.agrisera.com/) were used: RbcL AS03 037, PsaA AS06 172, PsbC (CP43) AS11 1787, AtpC AS08 312. NAB1 was kindly provided by Prof. Dr. Kruse form university of Bielefeld (Germany).

### Mitochondrial respiration

Samples in the exponential phase were subjected to respiratory rate measurements in the dark using a Clark-type O_2_ electrode (Oxygraph Plus; Hansatech Instruments; Clark, 1956). Respiratory rates were normalized to cells number obtained by Countless®II FL automated cell counter (Thermo Fisher). To discriminate between the individual contributions of the alternative and the cytochrome pathway dark respiration measurements were conducted as follows: cell samples (5*10^7^ cell/ml) were transferred to the measurement chamber of the Clark electrode, respiration rates were recorded for 3 min prior to the addition of the first inhibitor, then respiration rates were recorded for 3 additional min finally the second inhibitor was added and measurements were continued for another 3 min. Alternative respiration was inhibited by adding 2 mM SHAM (salicylhydroxamic acid), while the cytochrome pathway (complex III) was inhibited by adding 5µM myxothiazol. To assess the relative contribution of the cytochrome pathway, respiration was first measured in the absence of inhibitors (total dark respiration) before alternative respiration was inhibited by adding SHAM. Cytochrome dependent respiration was then inhibited using myxothiazol and the residual respiration determined in relation to the uninhibited state. The contribution of alternative respiration was determined by reversing the order of inhibitor addition (myxothiazol followed by SHAM) (Bailleul et al., 2015).

### NAB1 sequence analysis

*Chlamydomanas reinardtii* NAB1 protein (Cre06.g268600.t1.2) was blasted against *C. vulgaris* protein database (Cecchin et al., 2019) and *C. sorokiniana* protein database (https://greenhouse.lanl.gov/greenhouse/). Protein domains were obtained and aligned with Scanprosite tool of Prosite database (de Castro et al., 2006; Sigrist et al., 2013). Sequence alignment was visualized with Clustal Omega (Goujon et al., 2010; Sievers et al., 2011).

### Experimental replication and statistical treatment

All the experiments herein reported were performed at least three times. Errors are reported as standard deviations. Statistical significancy was tested by Tukey’s test.

## ACKNOWLEDGEMENTS

The research was supported by the ERC Starting Grant SOLENALGAE (679814) to M.B.. The lipid analyses were performed at the HelioBiotec platform (CEA Cadarache).

